# Cone mitochondria act as microlenses to enhance light delivery and confer Stiles-Crawford-like direction sensitivity

**DOI:** 10.1101/2021.02.11.430818

**Authors:** John M. Ball, Shan Chen, Wei Li

## Abstract

Evolution endeavors to maximize the function of biological structures in organisms, and the vertebrate eye is no exception. Cone photoreceptors in the retina are among the most energy-demanding cells in our body, necessitating numerous mitochondria. Intriguingly, these mitochondria adopt a peculiar spatial aggregation immediately beneath the cone outer segment (OS) that houses light-sensitive opsin molecules. Here we demonstrate, *via* direct live imaging and computational modeling of ground squirrel cones, that such mitochondria bundles concentrate light to enter the OS for detection. This “microlens”-like feature of cone mitochondria produces an angular dependence of light intensity quantitively consistent with the Stiles-Crawford effect, a psychophysical phenomenon believed to improve visual resolution. Thus, in addition to their function as a necessary powerhouse, cone mitochondria play a critical optical role.

## Main Text

Sensory systems that optimize the transduction of physical energy into biochemical signaling will have a significant evolutionary advantage. The vertebrate retina is a notable example, featuring an inverted structure with multiple neural layers through which photons must pass– risking premature absorption or scattering—prior to detection in distally located photoreceptor outer segments (OS) that contain light-sensitive opsin molecules. Rod photoreceptor nuclei (***1)*** and Müller glia (***2***) possess specialized optical features that may play roles in enhancing photon delivery. Birds and reptiles have evolved a fascinating array of colored oil droplets located in the distal extents of photoreceptor inner segments (IS), allowing fine-tuning of wavelength-dependent light sensitivity (***3–7***). Moreover, the established waveguiding of light by photoreceptors is instrumental to adaptive optics retinal imaging, which allows the observation of individual human cones using noninvasive methods (***8, 9***).

Mitochondria are essential intracellular organelles that provide energy for cellular functions. Although diverse in size, shape, number, and location in different cells and tissues, for most mammalian cells, they form a reticular network surrounding the nucleus (***10***). In the retina, however, photoceptors—especially cones—possess an abundance of mitochondria, tightly packed in into an elongated bundle (MtB) in many species including primates (***11***). In mammals, the MtB occupies the ellipsoid, the distal portion of the cone IS, immediately proximal to the OS (**Fig. 1A**). Such a high density of mitochondria at an unusual location is a peculiar feature of the retina, the exact biological value of which remains elusive.

**Figure 1:**
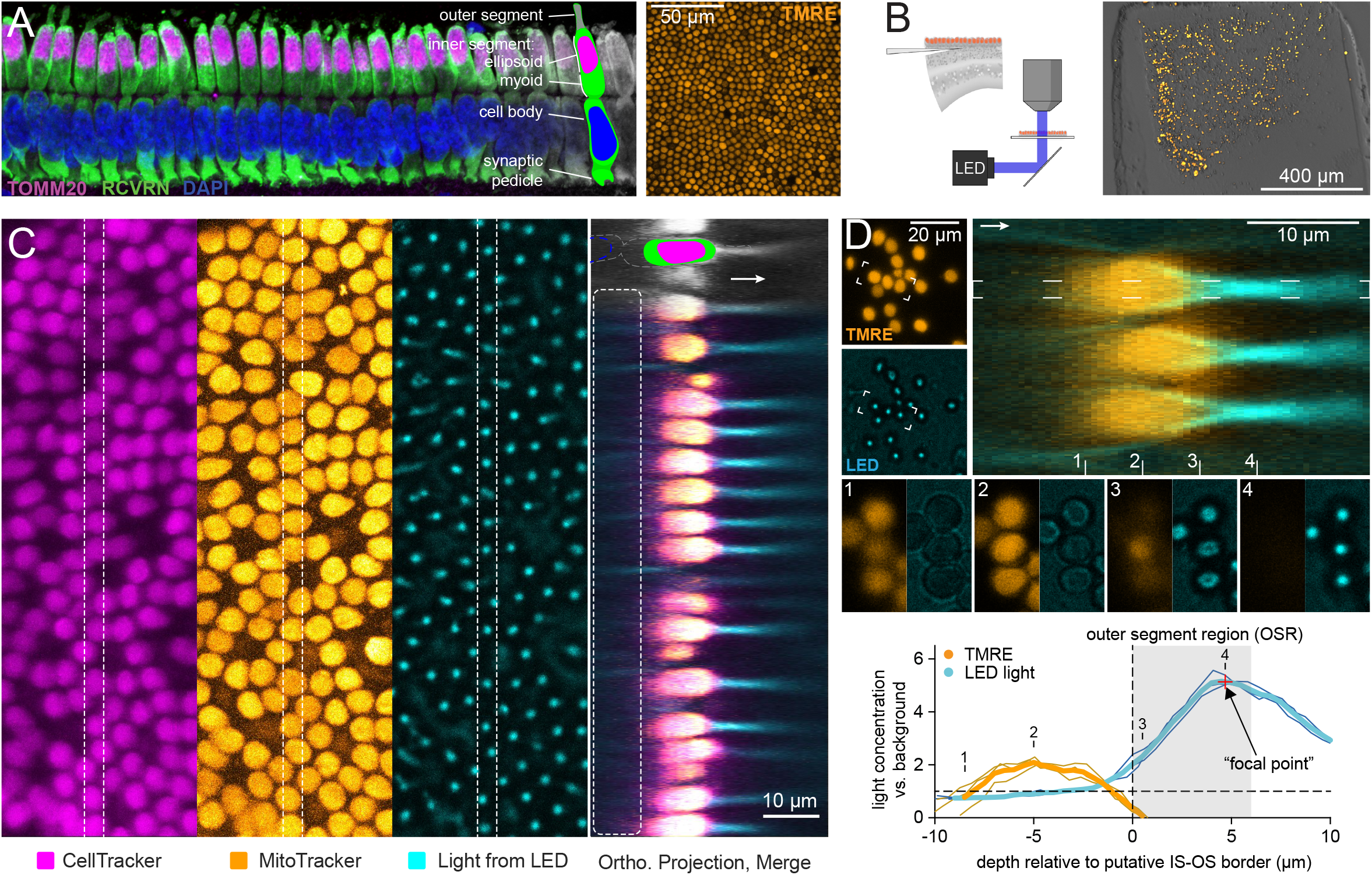
Light concentration by GS cone mitochondria. A) Cone mitochondria in superior GS retina. Left: Immunolabeled vertical section highlighting cone anatomy. Right: Live flat-mount view of TMRE-labeled mitochondria in photoreceptors. B) Horizontal sectioning and imaging of light transmission through agarose-embedded retinas. C) Concentration of light by photoreceptors in retinas sectioned as in ***B***. MitoTracker localizes to mitochondria; CellTracker labels cytoplasm. In the orthogonal projection, note the absence of CellTracker labeling where the cone myoid region would ordinarily be expected (dashed line; compare to ***A***). Arrow indicates the direction of light transmission. **D)** Quantification of light concentration factors in 3 exemplary cones. Flat-mount TMRE view is a maximum projection; LED image is a single plane at the peak concentration intensity. Orthogonal projection comes from the bracketed cones. Panels 1-4 depict features of light concentration at the depths indicated. Graph shows the relative light intensity (concentration factor) for the 3 cones (individual and average) as a function of depth from the distal IS tip in the 1.5 μm-diameter OS region (top cone, dashed line; see Methods). TMRE signal is arbitrarily scaled for comparison.

The undoubtedly sensible rationale for this configuration is to situate the MtB optimally to support the energy needs of the mighty molecular machinery of phototransduction in the OS (***12, 13***). However, it remains a mystery why mitochondria within the MtB require such a highly ordered spatial organization. Owing to their elevated index of refraction and conspicuous location—akin to the aforementioned oil droplets—it has been speculated that mitochondria may likewise play an important optical role (***11, 14, 15***). Nevertheless, due to the lack of a practical experimental preparation, a conclusive demonstration of their optical properties has yet to be presented.

To investigate the optics of the cone MtB, we took advantage of a unique species–the thirteen-lined ground squirrel (GS)–for its cone-dominant (85%) retina, featuring a single layer of photoreceptor cell bodies in its dorsal extent (**Fig. 1A**). Importantly, as a hibernator that endures near-freezing body temperatures for several months during the winter, the GS also features remarkable cold-tolerant cellular specializations that render *ex vivo* tissue stable for up to several days (***16–18***). These properties allowed us to develop a preparation in which the optical properties of live, mitochondria-packed cone ellipsoids could be observed. This preparation involved the isolation of the cone IS layer by horizontal vibratome-sectioning of agarose-embedded GS retinal explants (**Fig. 1B)**. The resulting samples featured patches of photoreceptors that often retained their native pseudo-hexagonal packing; their survival was verified by live labeling with tetramethylrhodamine ethyl ester (TMRE), which localizes to mitochondria due to their elevated inner membrane potential, providing both a clear indication of mitochondrial viability and a convenient marker for localization (**Fig. 1A-D**). Unexpectedly, such cones frequently lacked intact cell bodies, as assessed by post-hoc immunolabeling including DAPI (**fig. S1**) as well as during live imaging by Hoechst staining of cell nuclei (not shown) and the use of CellTracker, a fluorescent dye that permeates cell membranes and is subsequently converted to a cell-impermeant form (**Fig. 1C**). We speculate that the physical stress of sectioning may have caused photoreceptors to preferentially break at the cell body-IS junction (which is the narrowest part of the cell, normally supported by the outer limiting membrane), after which the plasma membrane sealed to form an ‘ellipsoidsome’ containing live mitochondria (**Fig. 1C** cartoon; **fig. S1A**). Further, cone OS were typically detached by the removal of the RPE, likely due to the tight ensheathment of OS by RPE interdigitation between cones, and their presence vs. absence did not impart any evident systematic differences upon the optical data presented below. Thus, although light absorption will certainly benefit further from the waveguiding properties of the OS itself, here we focus on the optical role of the cone ellipsoid in the delivery of light to the OS.

By replacing the condenser of a confocal microscope with a dichroic mirror to direct collimated light from a blue-filtered white LED (490nm cutoff) toward the sample (**Fig. 1B**), we were able to acquire 3D image stacks of TMRE fluorescence within MtB as well as a 3D snapshot of the distribution of light energy that traverses them (**Fig. 1C-D**). Strikingly, we observed that LED light passing through cones was concentrated into beams of light many times brighter than the background intensity (**Fig. 1D**). Near the ellipsoid base, weak “halos” of light were observed that inverted near the MtB midpoint and ultimately coalesced to bright spots (termed “focal points” here; **Fig. 1D**). These focal points were located a few mm beyond the distal end of the IS and had a half-width of ∼1 μm (**Fig. 1D**)—very similar to the size of the GS cone OS (∼6 μm long and 1.5 mm in diameter; see **Fig. 2A**). Such placement of a concentrated beam of light would greatly enhance entry of photons into the OS and thus, their detection. In fact, cones with shorter focal lengths would be better suited to couple this intense beam of light into the proximal end of the OS, which may then guide this light along its length, maximizing detection. Notably, in rare samples, sectioning yielded patches of isolated cones with intact cell bodies; cones in these patches featured short focal lengths (**fig. S2**), supporting the prediction that cones concentrate light at short distances *in vivo*.

**Figure 2:**
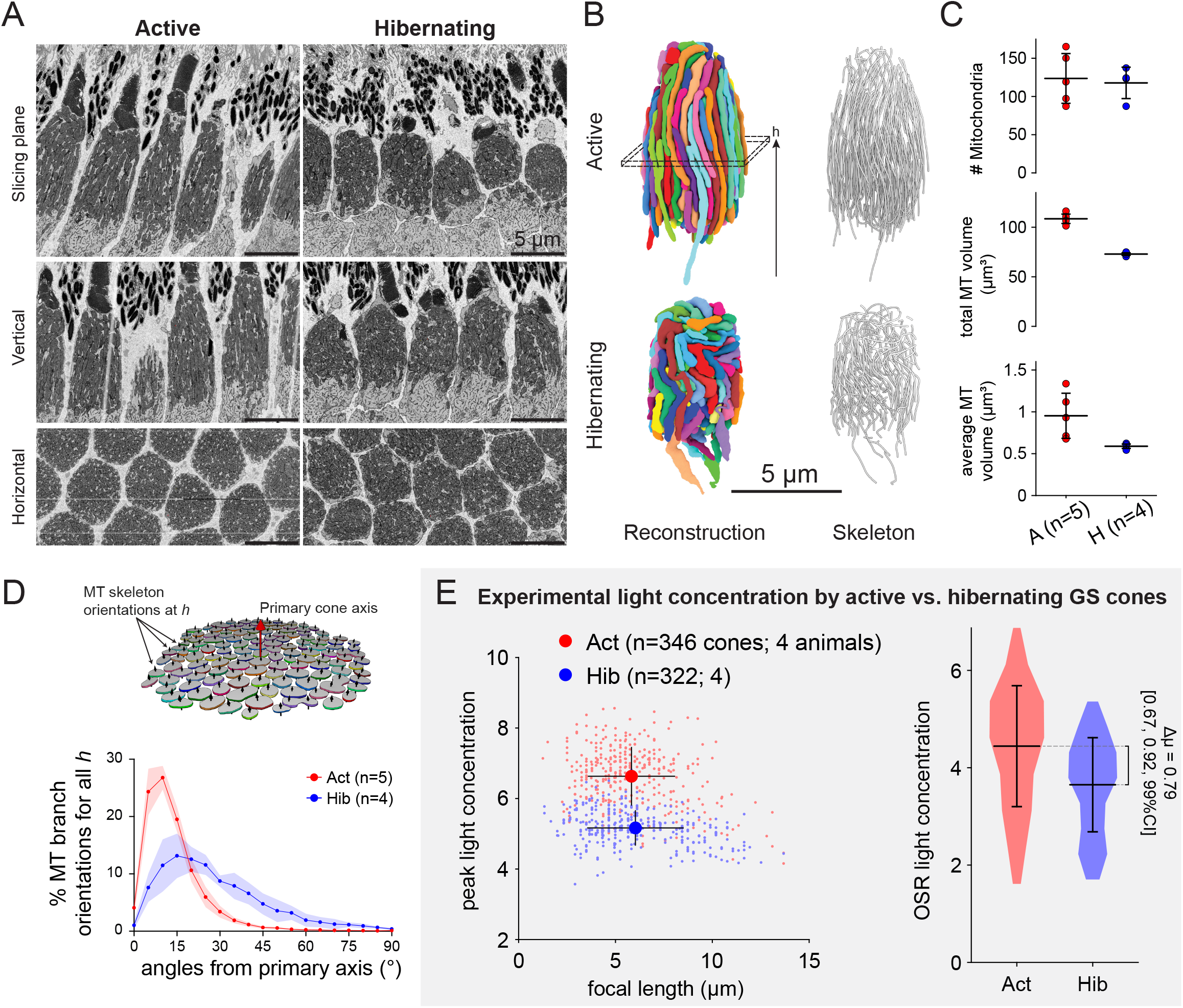
Cone mitochondrial structural changes correlate with hibernation-induced decreases in light gathering. A) SBEM images of GS cone mitochondria. Vertical and horizontal images are software projections. B) Example reconstructions (segmented 3D models and skeletonizations) of all mitochondria from sample cones. C) Morphological quantification of reconstructed mitochondria. Data shown as mean ± standard deviation. Statistical comparison not performed here (see Methods). D) Quantification of mitochondrial alignment in cones using skeletonizations. Diagram depicts the scheme for measuring mitochondria branch orientation at an example cone height *h* (see panel ***B***). Graph shows histograms (mean ± standard deviation; shaded regions) of mitochondria branch angles throughout reconstructed cones. **E)** Experimental light concentration comparison for active vs. hibernating GS cones (see **Fig. 1**). Scatterplot shows cone focal points and mean ± standard deviation for each condition. Violin plot shows the distributions of average light concentration in the OSR (see Methods for an explanation of statistics).

To further probe the relationship between MtB structure and this concentration of light, we took advantage of the hibernating GS, whose cone mitochondria have been reported to undergo structural changes that include a reduction in total number and/or volume (***19–21***). Such structural remodeling of the MtB may lead to measurable optical consequences. To obtain high quality 3D reconstructions of mitochondria, which yielded structures with a resolution suitable for computer simulation of light transmission (see details below), we applied serial block-face electron microscopy (SBEM) to dorsal cones in both an active and a hibernating GS (**Fig. 2A-B**). Mitochondria in hibernating GS cones were as numerous as those in awake GS but were individually smaller, resulting in a lower total MtB volume in hibernating cones (30% lower; see **Fig. 2B-C**). Importantly, mitochondria morphology in hibernating GS cones was qualitatively different; whereas mitochondria in active GS appeared elongated and well-organized, those in hibernating GS appeared distorted and markedly less well aligned (**Fig. 2B**). Detailed analysis indicated exceptional alignment among active GS cone mitochondria; approximately 75% of mitochondria skeleton “branches” reconstructed from awake cones deviated by less than 15° from overall MtB orientation (see Methods, **fig. S3** for details), compared to only 30% in hibernating cones, indicating considerable disorganization (**Fig. 2D; fig. S4**).

Concomitantly, we also observed optical differences between awake and hibernating GS cones. MtB from hibernating GS cones concentrated light less effectively than did those from active GS (**Fig. 2E**). While focal lengths were wide-ranging but similar in both awake and hibernating samples (active 5.8 ± 2.2 μm; hibernating 6.0 ± 2.5 μm), peak intensity was higher in active (6.6 ± 0.8-fold brighter than background) than in hibernating samples (5.2 ± 0.5-fold). In order to estimate the overall effect of these differences on the light theoretically available to a cone OS, we integrated the total light within a cylinder approximating the anatomical outer segment region (OSR; with dimensions as described above). Average light concentration in the OSR was ∼22% higher in active samples (4.44 ± 1.25 vs. 3.65 ± 1.0-fold-increase). This difference supports the hypothesis that the elongated, parallel organization of mitochondria in cone MtB appears to enhance the concentration of light for detection in the OS.

In such experiments, differences in MtB optics between active and hibernating GS may result from a confounding mixture of differences in gross morphology as well protein concentration (***22, 23***) or even altered cristae structure (***24***). To assess specifically the impact of MtB morphology on light concentration, we turned to computational methods. To this end, we translated reconstructed MtB models into 3D fine-grained dielectric structures (**Figs. 3A, S3**) suitable for use with code libraries from MEEP (MIT Electromagnetic Equation Propagation FDTD simulator), which divides time and space into a grid of points within which Maxwell’s equations of electromagnetism are calculated (***25***). The electromagnetic energy distribution in this grid, reached upon the introduction of a 450 nm continuous-wave energy source at the proximal end of the cone and allowing the resulting propagating waves to settle, provides a theoretical, 3D view of the passage of blue light through these structures that could be quantified with the same techniques used for our confocal imaging data (**Figs. 1-2**).

**Figure 3:**
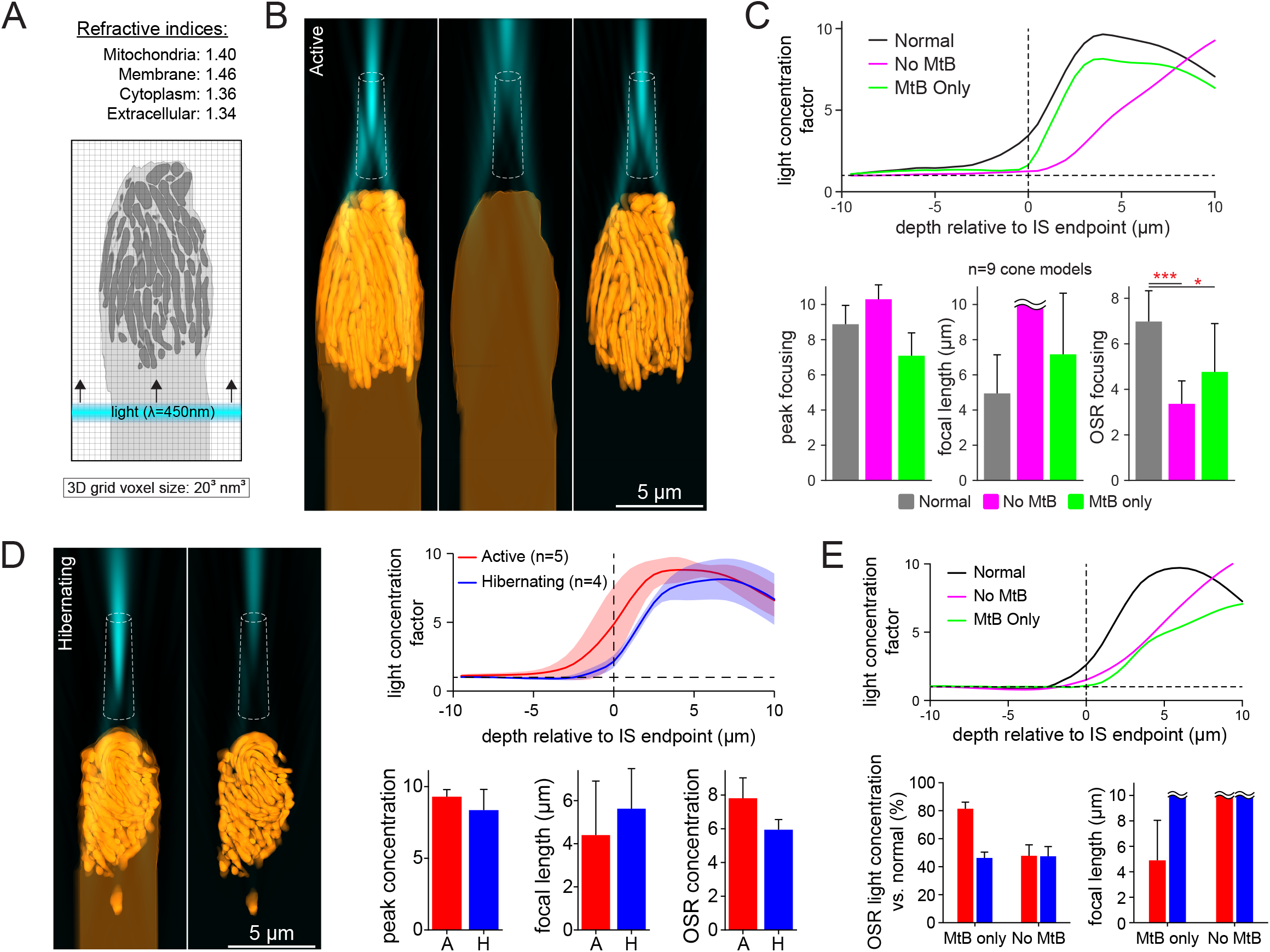
Electromagnetic simulations of light concentration by GS cone mitochondria. A) Simplified 2D schematic of the 3D FDTD simulation framework. B) Example light gathering by a reconstructed cone from active GS; from left-to-right: Intact, without mitochondria, suspended mitochondria only. Dashed shape depicts approximate OS region. C) Quantification of simulated light gathering. Top graph depicts light concentration factor profiles for simulations in ***B***. Bar graphs show aggregate measures across all reconstructions (n=9). Values are mean ± standard deviation; wavy lines indicate apparent focal lengths that fell outside the upper boundary of the simulation volume (> 10 μm). D) Simulated light gathering by an example cone from hibernating GS. Note the considerable decrease in intensity without the cell membrane (see also panel *E*). Graphs depict light gathering comparisons between active and hibernating GS cones (mean ± standard deviation). E) Mitochondria-dependence of simulated light gathering. Top profile plots are for simulations in ***D*** (left). Bottom graphs show change in concentration factor for active vs. hibernating GS cones with either the MtB or the IS cell membrane removed.

Light concentration by cones in simulations matched what we observed experimentally (**Fig. 3B**). Importantly, simulations permitted us to assess specifically the relative contribution of MtB structure and cone IS to the light concentration capability of cones. Cone IS without mitochondria were nonetheless capable of gathering light; however, in these simulations, concentration occurred at considerably longer focal lengths where the majority of the electromagnetic energy would fail to enter and be guided by or absorbed in the OS (**Fig. 3B-C**). In contrast, simulations of isolated MtB (as if the cone IS membrane had been dissolved, leaving behind only structurally intact mitochondria) exhibited light concentration much more similar to that of intact cones (**Fig. 3C**), suggesting that MtB play a major role in light concentration for entering OS.

Corroborating our experimental imaging results, simulations from hibernating GS cones likewise demonstrated weaker light concentration in the OSR (5.8 ± 0.6-fold vs. 7.8 ± 1.2-fold for active GS cones; **Fig. 3D**). This difference resulted both from lower peaks and longer focal distances -slightly differing from our experimental results, which largely reflect a reduction in peak intensity (**Fig. 2E**). This is likely due to other changes accompanying the structural alterations in mitochondria of hibernating GS cones, such as protein concentration (***26***) or inner matrix volume (***24***), which will affect mitochondrial refractive index, in turn altering focal lengths and peak intensity as demonstrated by simulations (**fig. S5**).

More importantly, isolated MtB from active GS cones were potent light gatherers, retaining 80% of the OSR light concentration factor seen in intact cone simulations (**Fig. 3E**); in contrast, isolated MtB from hibernating GS were only half as effective. Emphasizing this differential effect of MtB structure upon optics, cone IS devoid of mitochondria were equally ineffective at concentrating light, regardless of whether they came from active or hibernating GS (**Fig. 3E**). These results indicate that MtB with highly ordered spatial organization can significantly improve its ability to concentrate light. Indeed, simulations indicated that such organization was superior for light concentration compared to volume-matched globular mitochondria (**fig. S6**). Interestingly, a single “megamitochondrion” (**fig. S6**), an exaggerated configuration resembling the giant mitochondria seen in some cones of the tree shrew (***27***) and zebrafish (***28, 29***), similar also to the “ellipsosomes” present in other species (***30***), proved to be a highly effective configuration for light concentration.

Taken together, MtB configurations that minimize the refractive interfaces encountered by light by tight packing and elongation appear to maximize the potential for photon delivery to the OS. This feature, similar to the oil droplets encountered in birds and reptiles (***4***), evokes the notion of a microlens strategically placed for such a purpose. In fact, during imaging we observed that, like a biconvex lens, cone ellipsoids reproduced and inverted an image plane (generated by placing a mask with a slit in front of the LED) near their focal point (**Fig. 4A**). This observation likewise suggests that as with a lens, light incident upon MtB at an angle that deviates slightly from its optical axis may be nevertheless focused along that misaligned axis. In well-preserved retinal sections, alignment of inner and outer segments is typical, and may be an integral feature of light collection (**Fig. 4B**; see also ***31–33***). Therefore, light arriving along this axis will be maximally concentrated onto the OS; however, light entering at an angle will partially miss the OS (**Fig. 4B**). This observation could potentially contribute to the Stiles-Crawford effect (SCE), a psychophysical phenomenon rooted in photoreceptor optics that has captivated vision scientists for nearly 90 years (***34, 35***). The SCE, which dictates that photoreceptors (especially cones) are less sensitive to rays of light entering the pupil peripherally, is believed to improve visual resolution and reduce veiling background (**Fig 4C**; ***34***). Such directional sensitivity of light perception is canonically attributed to the waveguiding properties of cones (***35***). However, while the relatively high refractive index of mitochondria has been noted (***11, 36–38***), their role in SCE remains speculative, ranging from confounding the SCE via scattering (***36, 39***) to instead enhancing it (***11***). Here we hypothesize that coaxial light concentration by the MtB could produce SCE-like direction sensitivity, and the amount of light rendered unavailable to the OS due to misalignment would be affected by the extent of MtB focusing (**Fig. 4B**).

**Figure 4:**
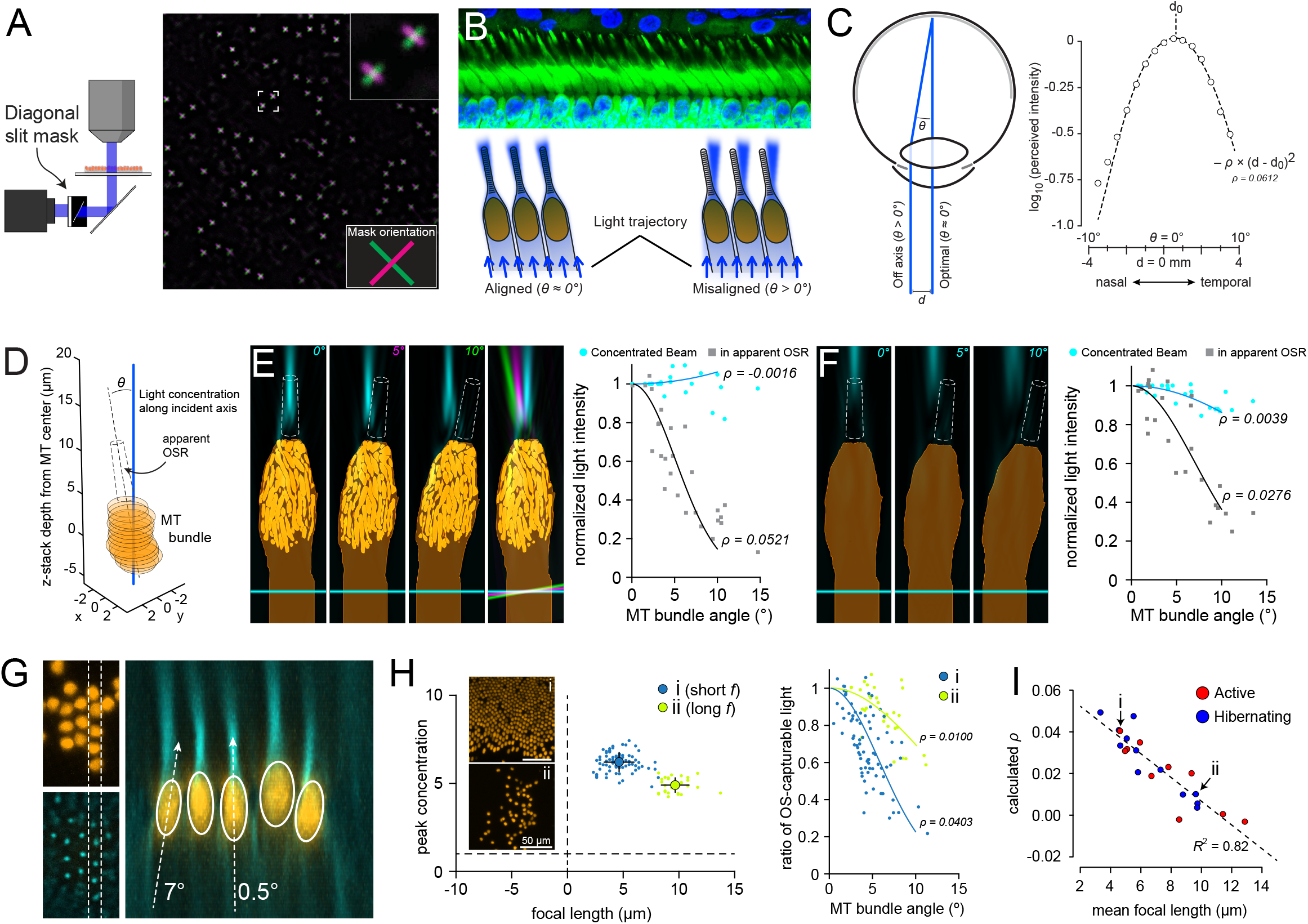
Stiles-Crawford-like direction-dependent light concentration by GS cone mitochondria. A) Reproduction of an inverted image plane (slit mask) at the focal points of cone MtB. Image is the overlaid result of two separate acquisitions with the mask reversed before the second acquisition. B) Conceptual illustration of angle-dependent concentration of light upon the cone OS. C) Diagram of the Stiles-Crawford effect and example two-sided quantification; reproduced from (***41***). D) Schematic of the quantification of direction sensitivity for cone light concentration in 3D image stacks (see Methods). E) Direction sensitivity of OSR light intensity in simulated GS cones. Left panels show example simulations with models tilted by the indicated angles. Overlay shows the simulations rotated in register with one another in different colors for comparison. Graph shows the direction sensitivity fit analysis (compare to panel ***C***) across simulations for all cone models (3 angles per cone model). F) Direction sensitivity of simulated cone light concentration without mitochondria. G) Concentration by cone mitochondria along the axis of light incidence. Annotations are simplified angle measurements in the 2D plane used here only for illustration. H) Increased direction sensitivity of OSR light concentration by cones in a sample with short average focal lengths (*i*; short *f*) compared to one with long average focal lengths (*ii*; long *f*). I) A strong correlation between mean focal length and apparent direction sensitivity (calculated *ρ*) across GS samples.

To test this, we simulated light transmission through GS cone models tilted by angles up to 10° and measured the energy density of light concentrated along the incident axis as well as in the apparent region in which the OS would reside if it were aligned with the tilting of the IS (**Fig. 4D-E**). We calculated the direction sensitivity parameter *ρ*, which originates from a fit to psychophysical SCE direction sensitivity data and describes the steepness of the SCE curve, and which ranges ∼0.05-0.08 mm^-2^ for foveal cones (**Fig. 4C**; see also ***40, 34, 41***), while smaller values can indicate pathological disruption of photoreceptors (***42***). Indeed, tilting had little effect on the amplitude of light concentration; however, the loss of light available to the shifted OS region as a function of tilting angle agreed well with the SCE, yielding *ρ* = 0.052 across all cone models (**Fig. 4E**). With the longer “focal lengths” produced in the absence of the MtB (**see Fig. 3C**), the loss of light at the OSR due to tilting was diminished, effectively “impairing” direction sensitivity (*ρ* = 0.028; **Fig. 4F**).

We were able to corroborate these theoretical results experimentally. Due to preparation and handling in live-imaging experiments, photoreceptors displayed apparent MtB tilts of up to 15° (**Fig. 4G**). While concentrated light beams in these cells appeared only minimally disturbed by misalignment, the light intensity within the presumptive OS showed a similar direction sensitivity (**Fig. 4H**). Moreover, as with simulations, samples with shorter focal lengths (likely reflecting healthier, more intact conditions) yielded larger *ρ* values, which were close to our simulation value and within the range of those reported for human SCE (**Fig. 4H**). This relationship between focal length and *ρ* was linear and strongly correlated across both active and hibernating TLGS (*R*^2^ = 0.82, **Fig. 4I**).

This demonstration of a role for MtB in the SCE-like angular dependence of concentrated light delivery to the cone OS provides necessary conceptual refinement to the waveguiding properties of cones. Central foveal (foveolar) cones are believed to possess more sparsely packed mitochondria compared to those even immediately outside the most central fovea (***11, 43, 44***). Interestingly, SCE is also weaker for foveolar cones than those outside the central fovea (***45–48***), which are shorter, have larger diameters, and feature prominent GS cone-like mitochondria bundles (***11, 34, 44***). Foveolar cones may however benefit from their long and narrow IS, as our simulations indicate that simply lengthening the inner segment of the GS cone shortens its effective focal length (**fig. S7**). Nonetheless, GS-like cone morphology is common among vertebrates, appearing even in lampreys (***49***). Taken together, direction sensitivity in non-foveolar cones—as well as in non-primate species—may require the MtB-based optical phenomenon demonstrated here.

In sum, through live-imaging and simulation, we demonstrated that spatial organization of cone mitochondria allows the MtB to efficiently concentrate light to enhance IS-OS coupling, possibly by minimizing the number of refractive interfaces that light must traverse, thus reducing light scatter and maximizing total internal refraction toward the OS. This simple optic feature also endows MtBs with a lens-like “focusing” ability that can produce SCE-like angular dependence of light delivery to the OS in non-foveolar cones; additional waveguiding of light by cone OS may further augment such directional sensitivity. Such MtB-dependent nature of this angular dependence may potentially be a non-invasive measurement for the diagnosis of retinal degenerations, many of which entail mitochondria dysfunction (***42, 50***).

## Acknowledgments

This work was funded by the intramural research program of the NIH. We thank Gerald Westheimer and Johnny Tam for helpful discussions and critical review of this manuscript. The simulation work for this study utilized the computational resources of the NIH HPC Biowulf cluster (http://hpc.nih.gov); we would like to extend special thanks to Susan Chacko for considerable assistance configuring and maintaining the MEEP simulation code libraries that made this study possible. We would like to thank the diligent work of the NIH animal facility staff for year-round care of the TLGS colony and hibernaculum. Mitochondria reconstructions were performed by the persistent, careful work of Anurag Goel, Jennifer Du, and David Lu under the supervision of Talia Kaden.

